# Optimized breeding strategies to harness Genetic Resources with different performance levels

**DOI:** 10.1101/2019.12.20.885087

**Authors:** Antoine Allier, Simon Teyssèdre, Christina Lehermeier, Laurence Moreau, Alain Charcosset

**Author notes:** Corresponding authors: Allier Antoine, RAGT 2n, Statistical Genetics Unit, 12510 Druelle, France, Alain Charcosset, GQE - Le Moulon, INRA, Univ. Paris-Sud, CNRS, AgroParisTech, Université Paris-Saclay, 91190 Gif-sur-Yvette, France.

## Abstract

The narrow genetic base of elite germplasm compromises long-term genetic gain and increases the vulnerability to biotic and abiotic stresses in unpredictable environmental conditions. Therefore, an efficient strategy is required to broaden the genetic base of commercial breeding programs while not compromising short-term variety release. Optimal cross selection aims at identifying the optimal set of crosses that balances the expected genetic value and diversity. We propose to consider genomic selection and optimal cross selection to recurrently improve genetic resources (i.e. pre-breeding), to bridge the improved genetic resources with elites (i.e. bridging), and to manage introductions into the elite breeding population. Optimal cross selection is particularly adapted to jointly identify bridging, introduction and elite crosses to ensure an overall consistency of the genetic base broadening strategy. We compared simulated breeding programs introducing donors with different performance levels, directly or indirectly after bridging. We also evaluated the effect of the training set composition on the success of introductions. We observed that with recurrent introductions of improved donors, it is possible to maintain the genetic diversity and increase mid- and long-term performances with only limited penalty at short-term. Considering a bridging step yielded significantly higher mid- and long-term genetic gain when introducing low performing donors. The results also suggested to consider marker effects estimated with a broad training population including donor by elite and elite by elite progeny to identify bridging, introduction and elite crosses.

## INTRODUCTION

Modern breeding has been successful in exploiting crop diversity for genetic improvement. However, current yield increases may not be sufficient in view of rapid human population growth (Godfray *et al.* 2010). Moreover, modern intensive breeding practices have exploited a very limited fraction of the available crop diversity (Cooper *et al.* 2001; Reif *et al.* 2005). The narrow genetic base of elite germplasm compromises long-term genetic gain and increases the genetic vulnerability to unpredictable environmental conditions (McCouch *et al.* 2013). Efficient genetic diversity management is therefore required in breeding programs. This involves the efficient incorporation of new genetic variation and its conversion into short- and long-term genetic gain.

Among the possible sources of diversity, wild relatives, exotic germplasm accessions and landraces that predate modern breeding exhibit substantial genetic diversity. These *ex-situ* genetic resources are conserved worldwide in international gene banks and national collections. They provide a promising basis to improve crop productivity, crop resilience to biotic and abiotic stresses and crop nutritional quality (Salhuana and Pollak 2006; Wang *et al.* 2017). In case of traits determined by few genes of large effect, the favorable alleles can be identified and introgressed into elite germplasm following established marker-assisted backcross procedures (e.g. Charmet *et al.* 1999; Servin *et al.* 2004; Han *et al.* 2017). Such introgressions have been successful for mono- and oligogenic traits (e.g. earliness loci in maize, Simmonds 1979; Smith and Beavis 1996 and SUB1 gene in rice, Bailey-Serres *et al.* 2010). Introgressions also proved to be successful for more polygenic traits where few major causal regions have been identified. For instance, Ribaut and Ragot (2006) successfully introgressed five regions associated with maize flowering time and yield components under drought conditions. For complex traits controlled by numerous genes with small effect, e.g. grain yield in optimal conditions, the identification and introgression of favorable alleles into elite germplasm were mostly unsuccessful (Bouchez *et al.* 2002). This requires to go beyond the introgression of few identified favorable alleles toward the polygenic enrichment of elite germplasm (Simmonds 1962, 1993). Although plant breeders recognize the importance of genetic resources for elite genetic base broadening, only little use has been made of it (Glaszmann *et al.* 2010; Wang *et al.* 2017). The main reason is that breeding progress continues (Duvick 2005; Tadesse *et al.* 2019) and that breeders are reluctant to compromise elite germplasm with unadapted and unimproved genetic resources (Kannenberg and Falk 1995). Despite genetic resources carry novel favorable alleles that may counter balance their low genetic value by an increased genetic variance when crossed to elites (Longin and Reif 2014; Allier *et al.* 2019b), their progeny performance is mostly insufficient for breeders. Thus, breeding strategies are needed to bridge the performance gap between genetic resources and elites and to transfer beneficial genetic variations into elite germplasm while not compromising the performance of released varieties (Simmonds 1993; Gorjanc *et al.* 2016). Pre-breeding can be defined as the recurrent improvement of genetic resources to release donors that can be further introduced into the elite breeding population (Figure 1). According to Simmonds (1993), pre-breeding should start from a broad germplasm and should be carried out on several generations with low selection intensity to favor extensive recombination events and minimal inbreeding. The donors released from pre-breeding can be directly introduced into the elite breeding population. However, in cases where the performance gap between the donors released from pre-breeding and elites is too large, one may consider a buffer population between donors and elites before introduction in the elite breeding population, further referred to as bridging. The best progeny of bridging is then considered for introduction into the elite breeding population (Figure 1).

**Figure 1.**
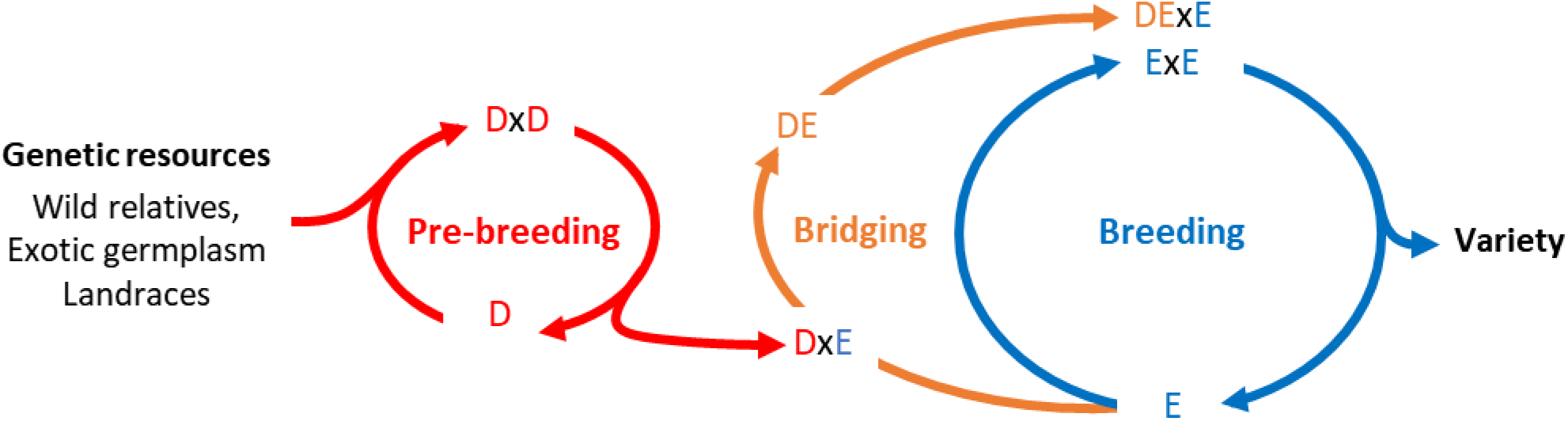
Diagram illustrating the respective positioning of pre-breeding, bridging and breeding from genetic resources to variety release.

Different sources of donors can be considered for genetic base broadening. This includes landraces historically cultivated before modern breeding. For instance in maize, open pollinated varieties (OPVs) are landrace populations of heterozygous individuals cultivated before the hybrid maize breeding revolution in the 1950’s (Anderson 1944; Troyer 1999). Inbred lines derived from OPVs present a large diversity and a potential interest for adaptation, but also a large performance gap with current varieties (Böhm *et al.* 2014; Melchinger *et al.* 2017; Böhm *et al.* 2017). These landraces can be further improved through pre-breeding that can be shared between the industry and public institutes in collaborative projects. In maize, the Latin American Maize Project (LAMP, Pollak 1990; Salhuana *et al.* 1997; Salhuana and Pollak 2006) provided breeders with useful characterization and evaluation of US and Latin American tropical germplasm accessions. Later, the Germplasm Enhancement of Maize project (GEM, Pollak and Salhuana 2001) improved the accessions identified in LAMP with elite lines furnished by private partners (Pollak 2003). Similarly, the Seed of Discovery project (SeeD, Gorjanc *et al.* 2016) aimed to harness favorable variations from landraces and to develop a bridging germplasm useful for genetic base broadening of commercial maize breeding programs. In this vein, Cramer and Kannenberg (1992) proposed the Hierarchical Open-ended Population Enrichment (HOPE) breeding system to release enriched maize inbreds for the industry. In its last version, the HOPE system is a breeding program with three hierarchical open ended gene pools permitting the transfer of favorable alleles from genetic resources to the elite pools (Popi 1997; Kannenberg 2001). Finally, breeders can consider the varieties released by breeding programs selecting on a different germplasm and in different environments as donors. In species where hybrid varieties are cultivated, the ability to use one variety’s inbred parent as a donor depends on the germplasm proprietary protection relative to species and countries (e.g. the possibility of using reverse breeding, Smith *et al.* 2008). In the US, maize inbred parents of hybrid varieties become publically available after twenty years of plant variety protection act, these are referred to as ex-PVPA (Mikel and Dudley 2006). In inbred species such as wheat, using current varieties for breeding is straightforward if cultivated under the union for the protection of new varieties of plants convention (UPOV, Dutfield 2011). These donors are likely the most performing but also the less original that can be considered.

With the availability of cheap high density genotyping, Whittaker *et al.* (2000) and Meuwissen *et al.* (2001) have proposed to use genomewide prediction to fasten breeding progress by shortening generation intervals. In the most frequently used approaches of genomewide prediction, it is assumed that most genomic regions equally contribute with relatively small effects to polygenic traits. A large number of genomewide markers is employed, and their effects are estimated on a training set (TS) of phenotyped and genotyped individuals. The genomic estimated breeding values (GEBVs) are further predicted considering the estimated marker effects and individuals’ molecular marker information. Recurrent selection based on genomewide prediction, further referred to as genomic selection (GS), has been increasingly implemented in crop breeding programs (Heslot *et al.* 2015; Voss-Fels *et al.* 2019). GS efficiency depends on the relationship between individuals in the TS and the target population of individuals to predict (Habier *et al.* 2010; Pszczola *et al.* 2012). As a consequence, in commercial breeding programs, GS has been mostly implemented considering a narrow elite TS that optimizes the prediction accuracy on elite material. However, such a narrow TS limits the prediction accuracy of individuals carrying rare alleles, which is the case for the progeny of elite by donor crosses. Therefore, it is important to define the TS composition that maximizes the prediction accuracy in both elite and introduction families.

In the context of genetic base broadening, GS is also interesting to fasten and reduce the costs for the evaluation and identification of genetic resources in gene banks (Crossa *et al.* 2016; Yu *et al.* 2016). Furthermore, GS can fasten pre-breeding programs to reduce the performance gap between genetic resources and elite populations (Gorjanc *et al.* 2016). Instead of truncated selection (i.e. select and mate individuals with the largest estimated breeding values), Cowling *et al.* (2017) proposed to use the optimal contribution selection to improve genetic resources while maintaining a certain level of diversity in the pre-breeding population. Optimal contribution selection (Wray and Goddard 1994; Meuwissen 1997; Woolliams *et al.* 2015) aims at identifying the optimal parental contributions to the next generation in order to maximize the expected genetic value in the progeny under a certain constraint on diversity. Therefore, the optimal contribution selection is particularly adapted to pre-breeding and genetic diversity management. Cowling *et al.* (2017) considered the pedigree relationship information but genomic relationship information can further improve the optimal cross selection (Clark *et al.* 2013). Considering optimal contribution selection on empirical cattle data, Eynard *et al.* (2018) observed that allowing for the introductions of old individuals in the breeding population increased long‐term response to selection. The optimal cross selection (OCS) is the extension of optimal contribution selection to deliver a crossing plan (Kinghorn *et al.* 2009; Kinghorn 2011; Akdemir and Isidro-Sánchez 2016; Gorjanc *et al.* 2018; Akdemir *et al.* 2019).

In this study, we propose to take advantage of OCS for selection of bridging, introduction and elite crosses (Figure 1). Allier *et al.* (2019d) proposed to account for within family variance and selection in a new version of OCS referred to as Usefulness Criterion Parental Contribution based OCS (UCPC based OCS). They observed both higher short- and long-term genetic gain compared to OCS in a simulated closed commercial breeding program. We extend here the use of UCPC based OCS to pre-breeding, following Cowling *et al.*(2017), and to an open commercial breeding program with recurrent introductions of genetic resources, extending the work of Eynard *et al.* (2018). Using OCS, the donor by elite crosses are selected complementarily to the elite by elite crosses in order to ensure an overall consistency of the genetic base broadening strategy. In this context, we aimed at evaluating the efficiency of genetic base broadening depending on the type of donors considered and the genetic base broadening scheme (Figure 1). We considered either donors corresponding to the generation of the founders of breeding pools or improved varieties released twenty years ago and five years ago. Our objectives were to evaluate (i) the interest of recurrent introductions of diversity in the breeding population, (ii) the interest to conduct or not bridging and (iii) the impact of the training set composition on within family genomewide prediction accuracies.

## MATERIAL AND METHOD

### Simulated breeding programs

#### Material and simulations

We considered 338 Dent maize genotypes from the Amaizing project (Rio *et al.* 2019; Allier *et al.* 2019c) as founders of genetic pools. This diversity was structured into three main groups: 82 Iowa Stiff Stalk Synthetics, 57 Iodents and 199 other dents. We sampled 1,000 biallelic quantitative trait loci (QTLs) with a minimal distance between two consecutive QTLs of 0.2 cM among the 40,478 single nucleotide polymorphisms (SNPs) from the Illumina MaizeSNP50 BeadChip (Ganal *et al.* 2011). Each QTL was assigned an additive effect sampled from a Gaussian distribution with a mean of zero and a variance of 0.05 and the favorable allele was attributed at random to one of the two SNP alleles. We sampled 2,000 SNPs as non-causal markers, further used as genotyping information. The consensus genetic positions of sampled QTLs and SNPs was considered according to Giraud *et al.* (2014).

We simulated two different breeding programs: an external breeding program (Figure 2A) that released every year varieties that were later considered as potential donors for introduction in a commercial breeding program (Figure 2C-D). Both external and commercial programs used doubled haploid (DH) technology to derive progeny. We assumed a period of three years to derive, genotype and phenotype DH progeny. Every year *T*, progeny of the three last generations *T*−3, *T*−4 and *T*−5 were considered as potential parents of the next generation. It created overlapping and connected generations as it can be encountered in breeding. We first considered a burn-in period of twenty years with recurrent phenotypic selection from a population of founders. Burn-in created extensive linkage disequilibrium as often observed in elite breeding programs (Van Inghelandt *et al.* 2011). Every progeny was phenotyped and phenotypes were simulated considering the genotypes at QTLs, an error variance corresponding to a trait repeatability of 0.4 in the founder population, and no genotype by environment interactions (File S1). Every individual was evaluated in four environments in one year. After twenty years of burn-in, we simulated different breeding programs using GS. Every year, progeny phenotypes and genotypes of the three last available generations were used to fit a G-BLUP model (File S1). Progeny were selected based on GEBVs and marker effects were obtained by back-solving the G-BLUP model (Wang *et al.* 2012) and further used for optimal cross selection to generate the next generation (see File S2).

**Figure 2.**
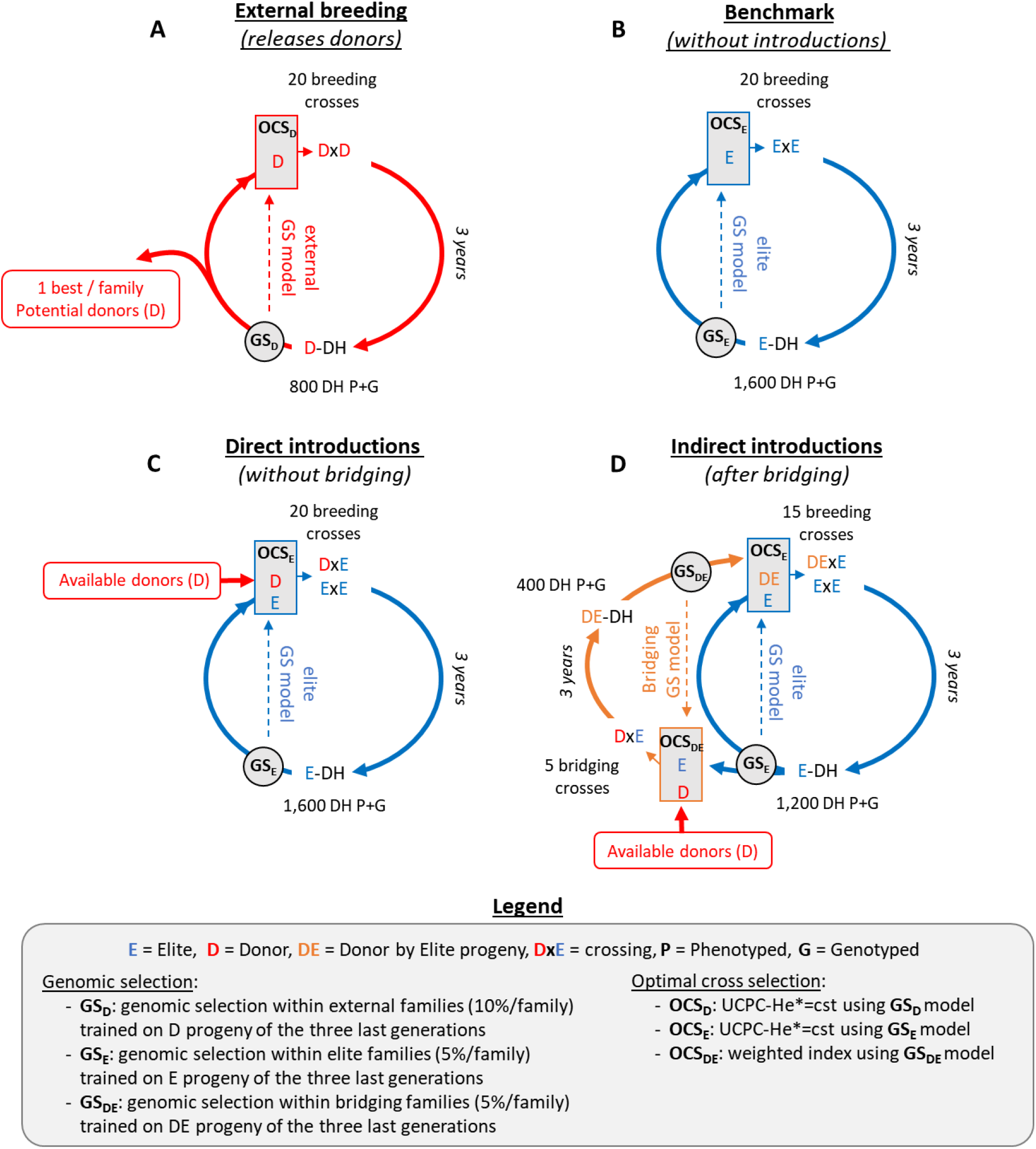
Diagram of simulated breeding programs. (A) External breeding program that generates potential donors, (B) commercial benchmark program without introductions, (C) commercial program with introductions without bridging or (D) commercial program with introductions after bridging.

#### External breeding program: Improvement of genetic resources

The external breeding program (Figure 2A) was simulated starting from a broad population of 40 founders sampled among the 338 maize genotypes. During the three first years, the founders were randomly crossed with replacement to generate each year 20 biparental families of 40 DH progeny to initiate the three overlapping generations. The genetic material in the external breeding is referred to as improved donors (D). During seventeen years, we first selected among the three last generations the 10% D progeny per family (i.e. 4 DH lines/family × 20 families × 3 years) with the largest phenotypic mean. We further randomly mated with replacement the 50 DH with the largest phenotypic mean to generate 20 biparental families of 40 DH lines. After twenty years of burn-in, we considered GS trained on the D progeny of the three last generations (i.e. 2,400 D progeny, Figure 2A). Among these three last generations, we considered per family the 10% D progeny with the largest GEBVs as potential parents of the next generation, i.e. N_D_ = 4 DH lines/family × 20 families × 3 years = 240 potential parents. The 20 two-way crosses among the N_D_(N_D_−1)/2 = 28,680 candidate crosses were selected using optimal cross selection (see optimal cross selection section).

#### Commercial breeding programs

The commercial breeding program (Figure 2B-D) started from a population of 10 founders sampled among the 57 Iodent genotypes. During the first three years, the founders were randomly crossed with replacement to generate each year 10 biparental families of 80 DH progeny to initiate the three overlapping generations. The elite genetic material in the internal breeding is referred to as elite progeny (E). During seventeen years, we considered as potential parents of the next generation the 50 E progeny with the largest phenotypic mean from the three last generations, i.e. without applying a preliminary within family selection. These were randomly mated to generate 20 biparental families of 80 DH lines. After twenty years of burn-in, we considered GS and differentiated three different scenarios: the benchmark commercial breeding program without introductions (Figure 2B), the commercial breeding program with direct introductions without bridging (Figure 2C) or the commercial breeding program with introductions after bridging (Figure 2D).

In absence of introductions (*benchmark*), the E progeny were selected based on the elite GS model trained on E progeny of the three last generations (i.e. 4,800 E progeny, Figure 2B). The 5% E progeny with the largest GEBVs within each family (i.e. 4 DH) in the three last breeding generations were considered as potential parents. The 20 two-way crosses among the 28,680 candidate ExE elite crosses were defined using optimal cross selection (see next section).

For scenarios with introductions, we considered different sub-scenarios for the genetic base broadening scheme (i) including (*Bridging*) or not bridging (*Nobridging*) and (ii) different types of potential donors, to cover different possibilities in both hybrid and inbred species. We considered as potential donors either the 338 genotypes from the Amaizing project or the D progeny with the largest GEBVs per family released by the external breeding program (i.e. 1 DH/family/year, 20 potential donors released every year). The scenario using the 338 genotypes from the Amaizing panel for genetic base broadening was identified with the suffix *Panel*. For the donors released by the external breeding program, we considered two time constraints for the access to diversity. To mimic a situation close to that of the US maize ex-PVPA system (Mikel and Dudley 2006), we considered donors released 20 to 24 years before the current year (i.e. 5 years × 20 DH = 100 potential D) in scenarios with the suffix *20y*. To simulate a faster access to external diversity, as it would be the case in line breeding under UPOV convention (Dutfield 2011), we considered the donors released by the external breeding 5 to 9 years before the current year (i.e. 100 potential D) in scenarios with the suffix *5y*.

For scenarios without bridging (Figure 2C), the E candidate parents were selected every year among the 5% E progeny showing the largest GEBVs per family in the three last breeding generations resulting in N_E_ = 4 DH × 20 families × 3 years = 240 potential E parents. The E progeny were selected based on the elite GS model trained on E progeny of the three last generations (i.e. 4,800 E progeny, Figure 2C). The 20 breeding crosses among the 28,680 candidate ExE elite crosses and the candidate DxE introduction crosses were selected using optimal cross selection and the elite GS model (see next section). Note that there was no constraint on the proportion of ExE elite or DxE introduction crosses.

For scenarios with bridging (Figure 2D), the population was split into a bridging population of 5 families of 80 DH (i.e. 400 DE progeny) and a breeding population of 15 families of 80 DH (i.e. 1,200 E progeny). Every year, the 15 breeding crosses where selected among all possible ExE elite and DExE introduction crosses. The E candidate parents for breeding were selected among the 5% E progeny per family showing the largest GEBVs from the three last breeding generations, resulting in N_E_ = 4 DH/family × 15 family × 3 year = 180 potential E parents. The E progeny were selected based on the elite GS model trained on all E progeny of the three last generations (i.e. 3,600 E progeny, Figure 2D). The DE candidate parents for introduction in the breeding population were similarly selected among the three last bridging generations, resulting in N_DE_ = 4 DH/family × 5 families × 3 years = 60 potential DE parents. The DE progeny were selected based on the bridging GS model trained on all DE progeny of the three last generations (i.e. 1,200 DE progeny, Figure 2D). Among the N_E_(N_E_ −1)/2 = 16,110 ExE possible elite crosses and the N_DE_N_E_= 10,800 DExE possible introduction crosses, 15 breeding crosses were selected using optimal cross selection with the elite GS model (see next section). Note that there was no constraint on the proportion of ExE elite or DExE introduction crosses. The 5 DxE bridging crosses were selected with the bridging GS model among the possible crosses between the available D and the E candidate parents conditionally to the 15 selected breeding crosses (see next section).

### Optimal cross selection

The optimal cross selection selects the set of crosses (***nc***) that maximizes the expected genetic value in the progeny (*V*) under a constraint on the genomewide genetic diversity in the progeny (*D*) (Kinghorn *et al.* 2009; Kinghorn 2011; Akdemir and Isidro-Sánchez 2016; Gorjanc *et al.* 2018; Akdemir *et al.* 2019). As proposed in Allier *et al.* (2019d), the effect of within family selection with intensity (*i*) and accuracy (*h*) on *V*^(*i*,*h*)^ and *D*^(*i*,*h*)^ can be accounted for in optimal cross selection by using UCPC based OCS (File S2). Similarly as in Allier *et al.* (2019d), we considered *h* = 1 for sake of simplicity.

For breeding crosses, the optimal set of |***nc***| = 20 crosses (in scenarios without bridging, Figure 2A-C) or |***nc***| = 15 crosses (in scenarios with bridging, Figure 2D) was selected to solve the multi-objective optimization problem:

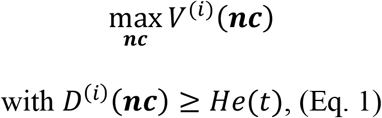

where *He*(*t*), ∀ *t* ∈ [0, *t*\*] is the minimal genomewide diversity constraint at time *t*. The evolution of diversity along time was controlled by the targeted diversity trajectory, i.e. *He*(*t*), ∀ *t* ∈ [0, *t*\*] where *t*\* ∈ ℕ* is the time horizon when the diversity *He*(*t*\*) = *He*\* should be reached. For the external and the commercial benchmark without introductions breeding programs, we considered *He*\* = 0.10 and *He*\* = 0.01 reached after sixty years, respectively. As in Allier *et al.* (2019d), the constraint on *D*^(*i*)^ followed a linear trajectory over time:

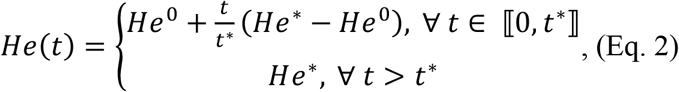

where *He*^0^ is the initial diversity at *t* = 0, i.e. at the end of burn-in.

For the commercial breeding program with introductions, we maintained the genomewide diversity constant after the end of burn-in, i.e. *He*(*t*) = *He*^0^, ∀ *t* ∈ ⟦0, *t*\*⟧. Thus, the UCPC based OCS selected introduction crosses (i.e. DxE if no bridging and DExE if bridging) when necessary to maximize the performance while keeping genomewide diversity constant (Eq. 1). In case of bridging, we completed the 15 selected breeding crosses with 5 bridging crosses (DxE, Figure 2D) that maximized the following function on the full set of |***nc***| = 20 crosses:

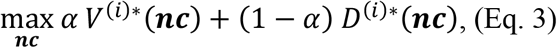

where, 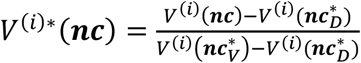 and 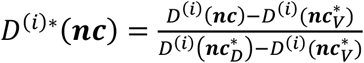 with 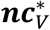 and 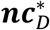 are the lists of crosses that maximize the performance (*V*) and the diversity (*D*), respectively, considering a within family selection intensity of *i*. *α* ∈ [0,1] is the relative weight given to performance compared to diversity. A differential evolution (DE) algorithm was used to find Pareto-optimal solutions of Eq. 1 and Eq. 3 (Storn and Price 1997; Kinghorn *et al.* 2009; Kinghorn 2011).

### Interest of pre-breeding and bridging

We compared different commercial breeding programs at a constant cost (i.e. total of 1,600 DH/year) with recurrent introductions (i) either direct or with a bridging step and (ii) considering three types of potential donors, resulting in the six genetic base broadening scenarios: *Bridging_Panel*, *Nobridging_Panel*, *Bridging_20y*, *Nobridging_20y*, *Bridging_5y*, *Nobridging_5y*. We ran ten independent simulation replicates of the external program that generated donors, the commercial benchmark without introductions, and the six genetic base broadening scenarios. Note that at a given simulation replicate the commercial breeding program accessed the potential donors released by the corresponding external breeding program simulation replicate.

We followed several indicators in the breeding families (i.e. E progeny, Figure 2). At each generation *T* ∈ [0,60] with *T* = 0 corresponding to the last burn-in generation, we computed the mean true breeding value (TBV) of E progeny *μ*(*T*) = *mean*(*TBV*(*T*)) and of the ten most performing E progeny 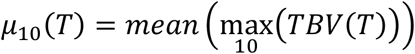 as a proxy of the performance that could be achieved at the commercial level by releasing these lines as varieties. We also measured the frequency of the favorable allele in the E progeny *p*_*j*_(*T*) at each QTL *j* among the 1,000 QTLs. We further focused on the QTLs where the favorable allele was rare at the end of burn-in, i.e. *p*_*j*_(0) ≤ 0.05. The results were averaged and standard errors were computed over ten independent replicates.

### Effect of a joint genomic selection model for bridging and breeding

For the three scenarios with bridging, we investigated the interest of a single TS grouping 3,600 DE and 1,200 E progeny to predict both breeding and bridging families. These three additional scenarios were referred to as *Bridging_Panel (Single TS)*, *Bridging_20y (Single TS)* and *Bridging_5y (Single TS)*. Every generation, we defined the prediction accuracies as the correlation between true breeding values and GEBVs (*cov*(*u, û*)) within breeding elite families (ExE), breeding introduction families (DExE) and bridging families (DxE). The prediction accuracies were averaged over the ten replicates and further averaged over the sixty generations. Note that considering a single GS model at constant cost yielded not only a broader but also a larger training set (4,800 DH progeny instead of 3,600 DH progeny for elite GS or 1,200 DH progeny for bridging GS, Figure 2).

We further investigated the effect of the proportion of DE and E progeny in the TS at constant size on within ExE and DExE family selection accuracy. We considered the 1,200 DE and 3,600 E progeny genotypes and phenotypes simulated at generations 18, 19, 20 in the first replicate of scenario *Bridging_20y*. We further selected the 5% DH per family with the highest GEBVs obtained using a GS model trained on all 4,800 progeny genotypes and phenotypes. These were randomly crossed to generate 50 elite (ExE) and 50 introduction (DExE) families of 80 DH progeny. These families were considered as the validation set (VS). We randomly sampled among the 4,800 DH progeny different TS of variable sizes and compositions (Table 1) and we evaluated the within elite (ExE) and introduction (DExE) family prediction accuracy (*cov*(*u, û*)). We also evaluated the within family variance prediction accuracy as the correlation between the variance of true breeding values and the estimated variance 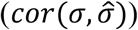. We reported results for twenty independent samples.

**Table 1.** Description of the training sets compared: the full training sets considering all available progeny of the last three generations and training sets at constant size (1,200 progeny or 3,600 progeny) with variable proportion of DE progeny.

|  | TS name | Number of E | Number of DE |
| --- | --- | --- | --- |
| **Full TS** | Pure E (3,600) | 3,600 | 0 |
|  | Pure DE (1,200) | 0 | 1,200 |
|  | 1/4 - DE (4,800) | 3,600 | 1,200 |
| **Constant size**<br>**(1,200)** | Pure E (1,200) | 1,200 | 0 |
|  | 1/4 - DE (1,200) | 900 | 300 |
| **Constant size**<br>**(3,600)** | 1/3 - DE (3,600) | 2,400 | 1,200 |
|  | 1/4 - DE (3,600) | 2,700 | 900 |
|  | 1/6 - DE (3,600) | 3,000 | 600 |
|  | 1/12 - DE (3,600) | 3,300 | 300 |
|  | 1/24 - DE (3,600) | 3,450 | 150 |
|  | 1/36 - DE (3,600) | 3,500 | 100 |

### Data availability statement

Supplemental files available at FigShare. File S1 contains additional material on the simulation of genotypes, the simulation of phenotypes and the genomewide prediction model considered. File S2 details the usefulness criterion parental contributions based optimal cross selection (UCPC based OCS). Supplemental Tables contains the supplemental tables S1-S3. Supplemental Figures contains the supplemental figures S1-S4. Data used in this manuscript are publically available at https://doi.org/10.25387/g3.7405892 and the R code of key functions can be found at https://doi.org/10.3389/fgene.2019.01006.

## RESULTS

### Interest of pre-breeding and bridging

The interest of recurrent introductions in the commercial breeding program after or without bridging depended on the type of donor considered. Panel donors showed a large performance gap with the elites they were crossed to. This performance gap increased with advanced breeding generations (on average a true breeding value difference with elites increased from −15 and −104 trait units). Improved donors showed a lower performance gap with elites. Twenty-year old donors showed an intermediate performance gap with elites (on average −22 trait units) and five-year old donors showed a reduced performance gap with elites (on average −8 trait units).

Direct introductions of panel donors without bridging (*Nobridging_Panel*) penalized the breeding population mean performance (*μ*) at short-term (at five years, *μ* = 8.168 +/− 0.282 compared to 9.239 +/− 0.237 without introductions, Figure 3A, Table S1) and long-term (at sixty years, *μ* = 9.651 +/− 0.958 compared to 38.837 +/− 1.563 without introductions, Figure 3A, Table S1). When considering the mean performance of the ten best progeny (*μ*_10_), the short-term penalty was no more significant (at five years, *μ*_10_ = 15.802 +/− 0.341 compared to 15.746 +/− 0.391 without introductions, Figure 3B, Table S2) but the long-term penalty was still significant (at sixty years, *μ*_10_ = 29.767 +/− 1.108 compared to 39.567 +/− 1.571 without introductions, Figure 3B, Table S2). The introduction of panel donors after bridging (*Bridging_Panel*) did not significantly penalize the short-term mean performance of the breeding population (at five years, *μ* = 8.688 +/− 0.329 compared to 9.239 +/− 0.237 without introductions, Figure 3A, Table S1) and yielded significantly higher long-term performance (at sixty years, *μ* = 52.110 +/− 0.886 compared to 38.837 +/− 1.563 without introductions, Figure 3A, Table S1). When considering *μ*_10_, the short-term penalty was reduced (at five years, *μ*_10_ = 15.605 +/− 0.477 compared to 15.746 +/− 0.391 without introductions, Figure 3B, Table S2) and the long-term gain increased (at sixty years, *μ*_10_ = 61.763 +/− 1.298 compared to 39.567 +/− 1.571 without introductions, Figure 3B, Table S2).

**Figure 3.**
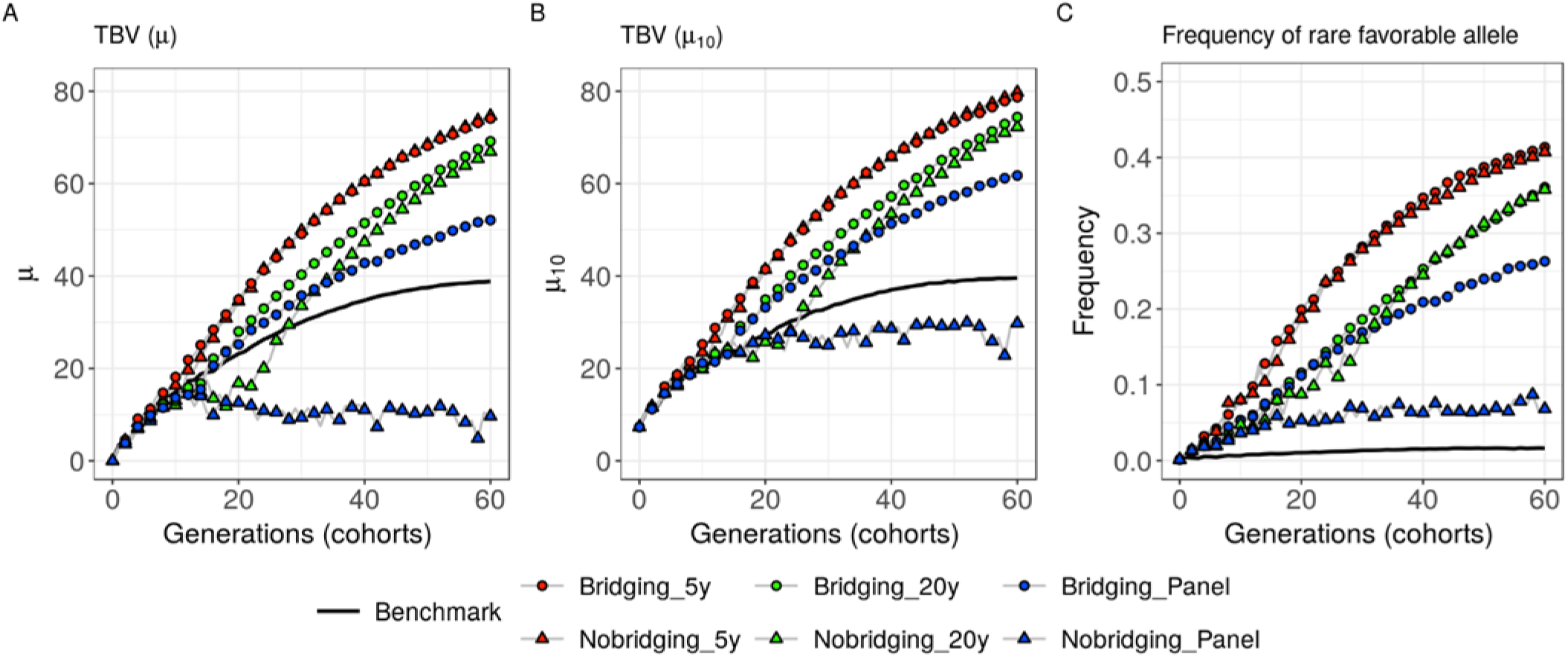
Evolution of the breeding population over generations. Scenarios considering presence or absence of bridging before introduction with different type of donors (panel, twenty-year old and five-year old donors). (A) Mean breeding population performance (*μ*), (B) mean performance of the ten best progeny (*μ*_10_) and (C) frequency of the favorable alleles that were rare at the end of burn-in (i.e. *p*(0) ≤ 0.05 corresponding on average to 269.9 +/− 23.6 QTLs).

Direct introductions of twenty-year donors without bridging (*Nobridging_20y*) yielded a penalty in the mid-term compared to not introducing donors (at twenty years, *μ* = 16.818 +/− 2.397 compared to 23.182 +/− 1.446 without introductions, Figure 3A, Table S1). When considering *μ*_10_, the mid-term penalty due to introductions was limited (Figure 3B, Table S2). After thirty years, this introduction scenario significantly outperformed the benchmark (*μ* = 33.546 +/− 1.519 compared to 30.006 +/− 1.319 without introductions, Figure 3A, Table S1) and this advantage increased until the end of the sixty years evaluated period (*μ* = 66.944 +/− 0.849 compared to 38.837 +/− 1.563 without introductions, Figure 3A, Table S1). The introduction of twenty-year old donors after bridging (*Bridging_20y*) penalized only the short-term performance (at five years, *μ* = 8.687 +/− 0.293 compared to 9.239 +/− 0.237 without introductions, Figure 3A, Table S1) and yielded significantly higher performance than the benchmark after twenty years (*μ* = 27.987 +/− 0.840 compared to 23.182 +/− 1.446 without introductions, Figure 3A, Table S1). Introductions after bridging significantly outperformed the direct introductions until the end of the sixty years evaluated period (*μ* = 69.154 +/− 0.868 with bridging compared to 66.944 +/− 0.849 without bridging and *μ*_10_ = 74.413 +/− 0.932 with bridging compared to 72.258 +/− 0.978 without bridging, Figure 3A-B, Table S1-S2).

Introducing five-year old donors after or without bridging yielded significantly higher mid- and long-term performances than all other tested scenarios, without any significant long-term advantage of introductions after bridging compared to direct introductions (at sixty years, *μ* = 74.074 +/− 0.869 with bridging compared to 74.662 +/− 0.938 without bridging, Figure 3, Table S1).

We observed that the recurrent introductions of donors impacted the genetic diversity of the commercial germplasm. The faster the commercial program had access to recent germplasm of the external program, the more the varieties released by the commercial program were admixed with the external program elite germplasm (Figure 4B and Figure 4C). In the scenario where only panel donors were accessible for introductions, the internal program diversity did not converge toward the external program (Figure 4A).

**Figure 4.**
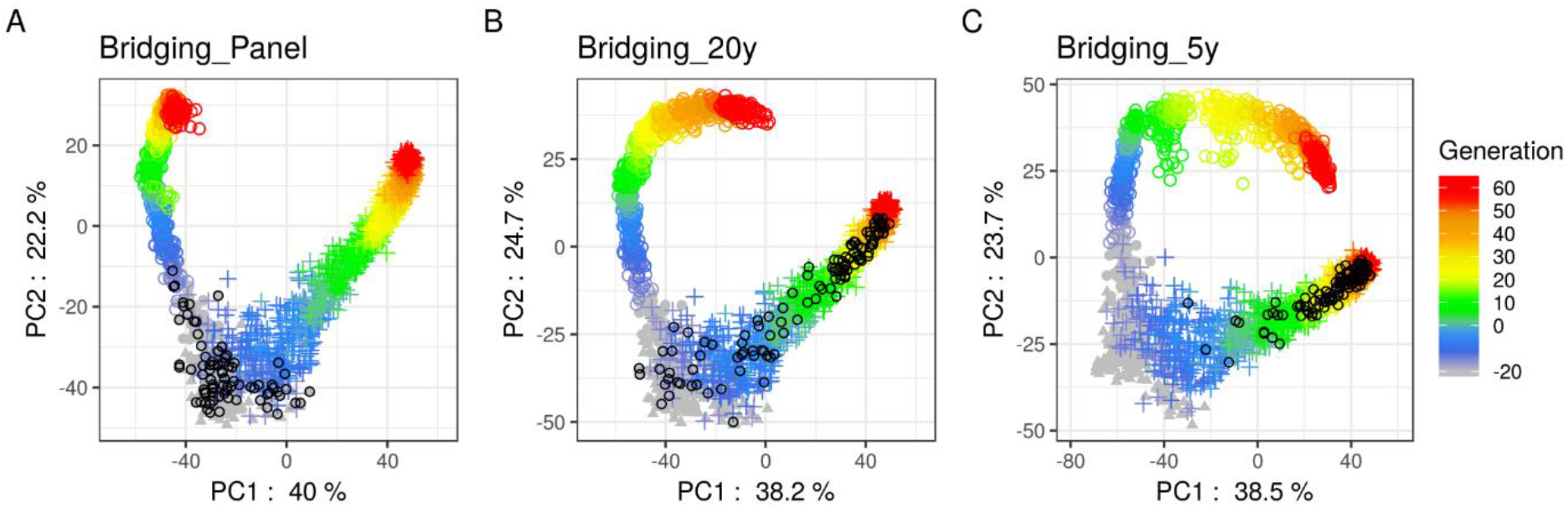
Principal component analysis of the modified Roger’s genetic distance matrix (Wright 1978) of the 338 founders (gray: points for the 57 Iodent lines and triangles for the 281 remaining lines), the commercial ten best performing E progeny per generation (colored circle sign) and the twenty donors per generation released by the external program (colored plus sign). Both commercial and external lines are colored regarding their generation (note that negative generations correspond to burn-in). Black circles represent the donors that have been introduced into the commercial breeding program. Only three scenarios with bridging are represented for the first simulation replicate, (**A**) when only donors from panel were accessible, (**B**) when twenty-year old donors from the external breeding were accessible and (**C**) when five-year old donors from the external breeding were accessible.

The evolution of the mean frequency of initially rare favorable alleles (i.e. favorable allele that had a frequency at the end of burn-in ≤ 0.05 in the elite breeding population) also highlighted differences between strategies. The older the donors, the lower the observed increase in frequency of initially rare favorable alleles (at sixty years for scenario with bridging, the mean frequency was 0.414 +/− 0.012 for five-year old donors, 0.361 +/− 0.009 for twenty-year old donors, 0.263 +/− 0.008 for panel donors and 0.016 +/− 0.006 without introductions, Figure 3C, Table S3). For twenty-year old donors, omitting the bridging before introduction delayed the increase in frequency of initially rare favorable alleles (e.g. at twenty years, the mean frequency was 0.088 +/− 0.014 without bridging compared to 0.116 +/− 0.011 with bridging, Figure 3C, Table S3). For panel donors the absence of bridging significantly penalized the increase in frequency of initially rare favorable alleles (at sixty years, 0.068 +/− 0.007 without bridging compared to 0.263 +/− 0.008 with bridging, Figure 3C, Table S3).

### Effect of a joint genomic selection model for bridging and breeding

Scenarios considering a single TS of 3,600 E and 1,200 DE progeny yielded higher mid- and long-term *μ* and *μ*_10_ than scenarios considering two distinct TS for bridging and breeding (Figure 5A-B). After twenty years, single TS scenarios significantly outperformed scenarios with two distinct TS (*μ* = 40.111 +/− 1.149 compared to 34.900 +/− 0.905 for five-year old donors, *μ* = 30.497 +/− 1.135 compared to 27.987 +/− 0.840 for twenty-year old donors and *μ* = 29.292 +/− 0.802 compared to 25.212 +/− 1.314 for panel donors, Figure 5A, Table S1). After sixty years, the advantage of a single TS remained significant except for five-year old donors (*μ* = 75.749 +/− 1.093 compared to 74.074 +/− 0.869 for five-year old donors, *μ* = 71.130 +/− 1.028 compared to 69.154 +/− 0.868 for twenty-year old donors and *μ* = 57.067 +/− 1.444 compared to 52.110 +/− 0.886 for panel donors, Figure 5A, Table S1). When considering *μ*_10_, a single TS was still more performing but its interest was less significant (e.g. for panel donors after sixty years, *μ*_10_ = 63.699 +/− 1.698 compared to 61.763 +/− 1.298, Figure 5 B, Table S1-S2). A single TS also favored the increase in frequency of initially rare favorable alleles introduced by five-year old donors and twenty-year old donors (e.g. for twenty-year old donors after sixty years, 0.380 +/− 0.010 compared to 0.361 +/− 0.009, Figure 5C, Table S3).

**Figure 5.**
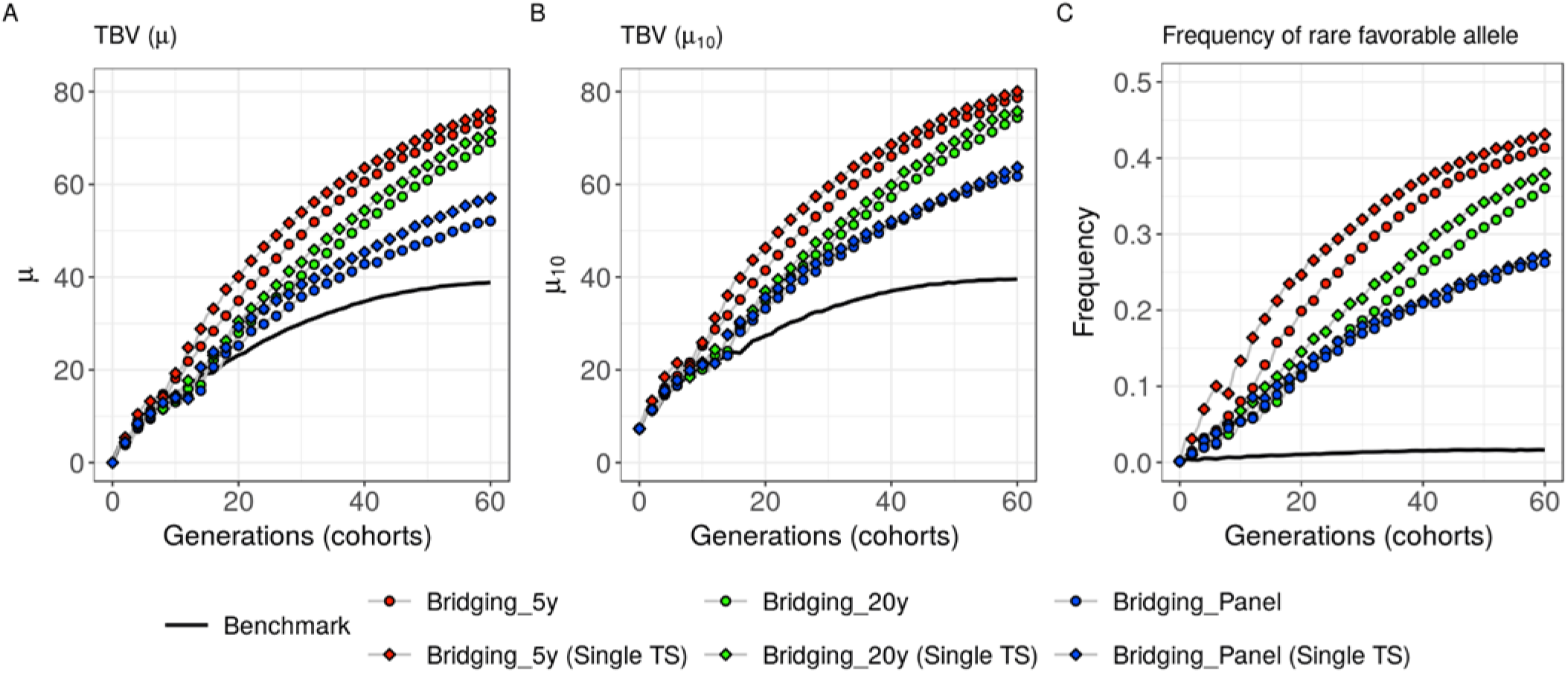
Evolution of the breeding population over generations. Scenarios considering bridging with different donors (panel, twenty-year old and five-year old donors) and either a single broad TS (*Single TS*) or two distinct training sets for bridging and breeding (*default*). (A) Mean breeding population performance (*μ*), (B) mean performance of the ten best progeny (*μ*_10_) and (C) frequency of the favorable alleles that were rare at the end of burn-in (i.e. *p*(0) ≤ 0.05 corresponding on average to 269.9 +/− 23.6 QTLs).

The observed within family prediction accuracies varied depending on the TS considered. For twenty-year old donors introduced after bridging, considering a single TS of 4,800 DE+E did not significantly improve the prediction accuracy within ExE families compared to using the pure elite TS of 3,600 E (*cor*(*u, û*) = 0.73 +/− 0.06 compared to *cor*(*u, û*) = 0.72 +/− 0.07, Table 2). However, it significantly improved the prediction accuracy within introduction DExE families compared to the pure elite TS of 3,600 E (*cor*(*u, û*) = 0.77 +/− 0.07 compared to *cor*(*u, û*) = 0.61 +/− 0.11, Table 2). A single TS also slightly but not significantly improved the prediction accuracy within bridging DxE families compared to the pure bridging TS of 1,200 DE (*cor*(*u, û*) = 0.78 +/− 0.05 compared to *cor*(*u, û*) = 0.73 +/− 0.06, Table 2). Similar observations were made on the other scenarios considering five-year old and panel donors. Prediction accuracies were larger in introduction DExE and bridging DxE families with older donors, i.e. phenotypically distant to elites, due to larger within family variances (e.g. for DExE families 14.43 +/− 4.40 for panel donors, 6.92 +/− 2.10 for twenty-year old donors and 5.00 +/− 1.41 for five-year old donors, Table 2).

**Table 2.** Within family prediction accuracies (*cor*(*u, û*)) depending on the validation set (VS): elite (ExE), introduction (DExE) and bridging (DxE) and the training set (TS) considered: pure elite (E), pure bridging (DE) and merged (E+DE). Results are given for scenarios with different donors, from the panel, twenty-year old and five-year old donors, considering a single TS and prediction accuracies are averaged over the ten replicates and all sixty generations. In brackets are given the standard errors averaged over sixty generations.

|  | Five-year old donor |  |  |  | Twenty-year old donor |  |  |  | Panel donor |  |  |  |
| --- | --- | --- | --- | --- | --- | --- | --- | --- | --- | --- | --- | --- |
|  | Family<br>variance | Prediction accuracy |  |  | Family<br>variance | Prediction accuracy |  |  | Family<br>variance | Prediction accuracy |  |  |
|  |  | TS = | TS = | TS = |  | TS = | TS = | TS = |  | TS = | TS = | TS = |
|  |  | E<br>(3,600) | DE<br>(1,200) | E+DE<br>(4,800) |  | E<br>(3,600) | DE<br>(1,200) | E+DE<br>(4,800) |  | E<br>(3,600) | DE<br>(1,200) | E+DE<br>(4,800) |
| VS =<br>**ExE** | 3.76<br>(1.17) | 0.69<sup>a</sup><br>(0.07) | 0.48<br>(0.1) | 0.72<sup>b</sup><br>(0.06) | 3.93<br>(1.06) | 0.72<sup>a</sup><br>(0.07) | 0.47<br>(0.10) | 0.73<sup>b</sup><br>(0.06) | 4.02<br>(1.16) | 0.72<sup>a</sup><br>(0.05) | 0.44<br>(0.10) | 0.73<sup>b</sup><br>(0.05) |
| VS =<br>**DExE** | 5.00<br>(1.41) | 0.60<sup>a</sup><br>(0.1) | 0.59<br>(0.1) | 0.73<sup>b</sup><br>(0.07) | 6.92<br>(2.10) | 0.61<sup>a</sup><br>(0.11) | 0.65<br>(0.10) | 0.77<sup>b</sup><br>(0.07) | 14.43<br>(4.40) | 0.65<sup>a</sup><br>(0.12) | 0.78<br>(0.07) | 0.86<sup>b</sup><br>(0.05) |
| VS =<br>**DxE** | 9.69<br>(2.01) | 0.61<br>(0.08) | 0.66<sup>a</sup><br>(0.08) | 0.73<sup>b</sup><br>(0.07) | 18.31<br>(3.78) | 0.65<br>(0.08) | 0.73<sup>a</sup><br>(0.06) | 0.78<sup>b</sup><br>(0.05) | 64.15<br>(12.89) | 0.74<br>(0.07) | 0.82<sup>a</sup><br>(0.04) | 0.86<sup>b</sup><br>(0.03) |

<sup>a</sup> Prediction accuracies that would have been realized if the breeding (E) or bridging (DE) set had been each predicted only by the corresponding training set (to be compared with <sup>b</sup>).
<sup>b</sup> Realized prediction accuracies when considering a single training set (to be compared with <sup>a</sup>).

At constant TS size of 3,600 DH, the increase in proportion of DE progeny from 0 to 1/3 in the TS increased the prediction accuracy within introduction DExE families (*cor*(*u, û*) = 0.58 +/− 0.02 to 0.73 +/− 0.01, Figure 6B) while it reduced the prediction accuracy within elite ExE families (*cor*(*u, û*) = 0.70 +/− 0.01 to 0.65 +/− 0.02, Figure 6A). The TS with 3,000 E and 600 DE appeared as a suitable compromise with within introduction DExE family *cor*(*u, û*) = 0.70 +/− 0.02 and elite ExE families *cor*(*u, û*) = 0.68 +/− 0.01. At constant TS size of 1,200 DH, the TS with 900 E and 300 DE progeny performed similarly as the pure bridging TS for prediction within DExE families (*cor*(*u, û*) = 0.63 +/− 0.03 compared to 0.62 +/− 0.02, Figure 6B) but significantly outperformed the pure bridging TS for prediction within elite ExE families (*cor*(*u, û*) = 0.52 +/− 0.04 compared to 0.34 +/− 0.02, Figure 6A). The within family variance prediction accuracy showed similar tendencies (Figure 7A-B). The increase in proportion of DE progeny from 0 to 1/3 in the TS increased the prediction accuracy within introduction DExE families (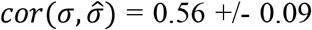 to 0.76 +/− 0.07, Figure 7B) while it slightly reduced the prediction accuracy within elite ExE families (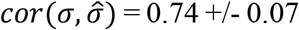 to 0.71 +/− 0.08, Figure 7A).

**Figure 6.**
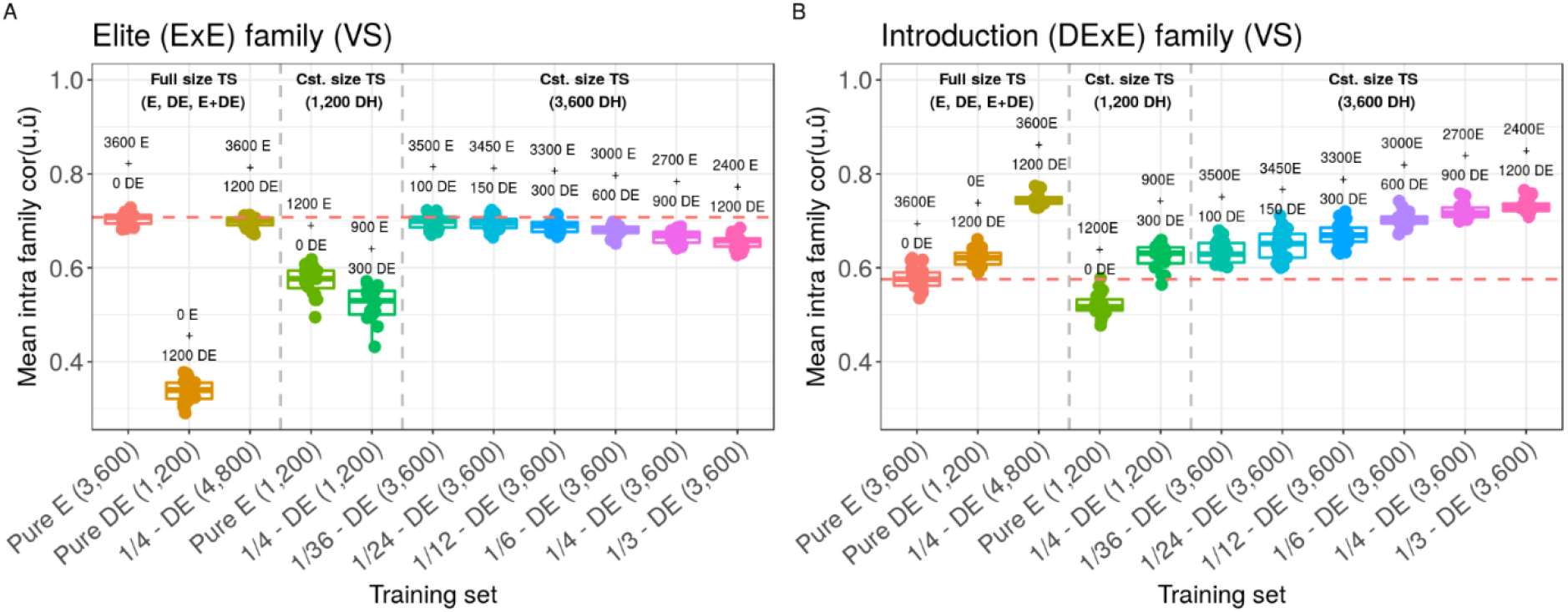
Effect of TS composition on intra family prediction accuracies (*cor*(*u, û*)) considering genotypes simulated at generations 18, 19, 20 in the scenario *Bridging_20y*. (A) Mean prediction accuracy within 50 elite (ExE) families and (B) mean prediction accuracy within 50 introduction (DExE) families. Boxplots represent the results for 20 independent replicates. One can distinguish three training set types (left to right): Full training set considering all 3,600 E progeny (Pure E), all 1,200 DE progeny (Pure DE) and all 3,600 E + 1,200 DE progeny; Training sets at constant size of 1,200 DH for comparison with Pure DE; Training sets at constant size of 3,600 DH and variable proportion of DE progeny for comparison with Pure E. The red dotted line represents the median value for Pure E TS.

**Figure 7.**
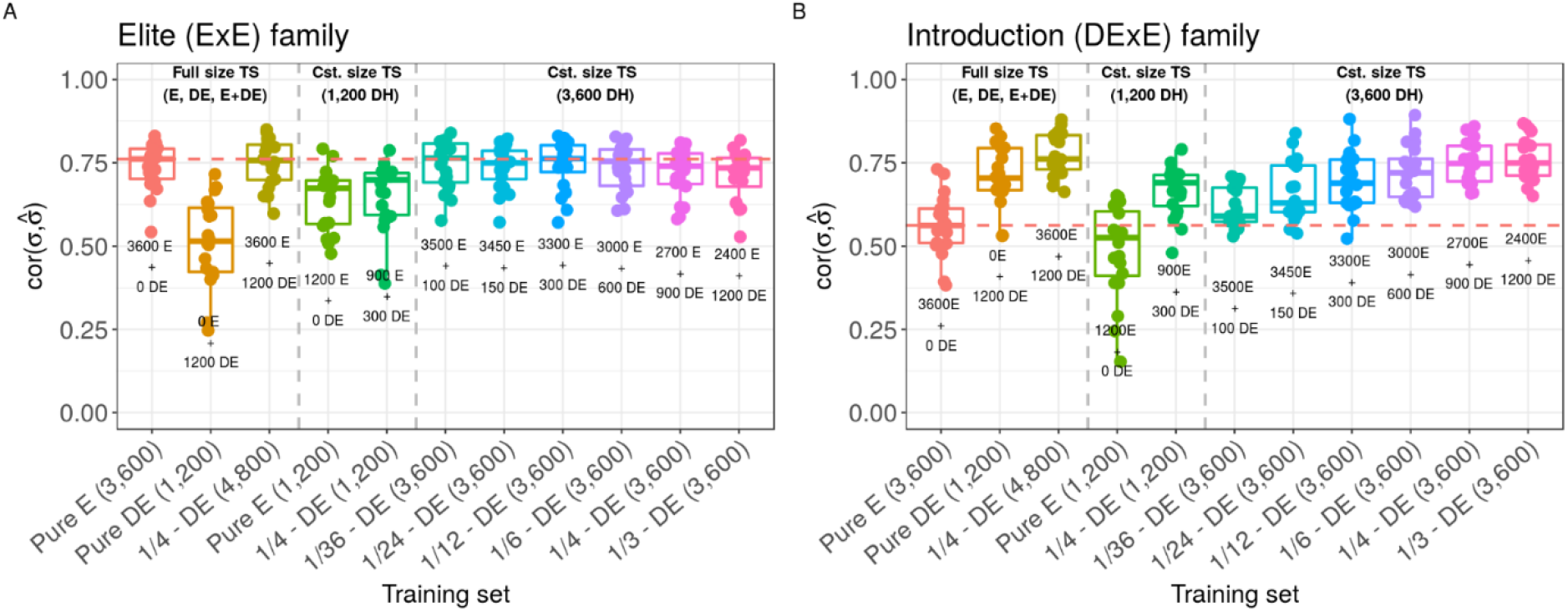
Effect of TS composition on family variance prediction accuracy 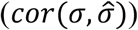 considering genotypes simulated at generations 18, 19, 20 in the scenario *Bridging_20y*. (A) Mean prediction accuracy in 50 elite (ExE) families and (B) mean prediction accuracy in 50 introduction after bridging (DExE) families. Boxplots represent the results for 20 independent replicates. One can distinguish three training set types (left to right): Full training set considering all 3,600 E progeny (Pure E), all 1,200 DE progeny (Pure DE) and all 3,600 E + 1,200 DE progeny; Training sets at constant size of 1,200 DH for comparison with Pure DE; Training sets at constant size of 3,600 DH and variable proportion of DE progeny for comparison with Pure E. The red dotted line represents the median value for Pure E TS.

## DISCUSSION

Despite the recognition of the importance to broaden the elite genetic base in most crops, commercial breeders are reluctant to penalize the result of several generations of intensive selection by crossing these to unimproved genetic resources. Furthermore, among the large diversity available for genetic base broadening (e.g. landraces, public lines, varieties…), the identification of the useful genetic diversity to broaden the elite pool is difficult and might dishearten breeders. Consequently, there is a need for global breeding strategies that identify interesting sources of diversity that complement at best the elite germplasm, improve genetic resources to bridge the performance gap with elites, and efficiently introduce them into elite germplasm.

### Genetic base broadening with optimal cross selection accounting for within family variance

The identification of genetic resources for polygenic enrichment of the elite pool should account for the complementarity between genetic resources and elites as reviewed in Allier *et al.* (2019c). Allier *et al.* (2019b) proposed the Usefulness Criterion Parental Contribution (UCPC) approach to predict the interest of crosses between genetic resources and elite recipients based on the expected performance and diversity in the most performing fraction of the progeny. The interest of UCPC relies on the fact that it accounts for within family variance and selection when identifying crosses. For instance, when crossing phenotypically distant parents, e.g. genetic resource and elite recipient, we expect a higher cross variance that should be accounted for to properly evaluate the usefulness of the cross (Schnell and Utz 1975; Longin and Reif 2014; Allier *et al.* 2019b). Additionally, we expect the best performing fraction of the progeny to be genetically closer to the best parent. This deviation from the average parental value should be considered to evaluate properly the genetic diversity in the next generation (Allier *et al.* 2019b; d). Accounting for parental complementarity at marker linked to QTLs also favors effective recombination in progeny and breaks negative gametic linkage disequilibrium between QTLs (i.e. repulsion), which unleashes additive genetic variance and increases long-term genetic gain (Allier *et al.* 2019d). Therefore, the OCS is particularly adapted to genetic diversity management in pre-breeding and breeding programs (Akdemir and Isidro-Sánchez 2016; Cowling *et al.* 2017; Gorjanc *et al.* 2018; Allier *et al.* 2019d). Based on these studies, we evaluated a UCPC based OCS strategy to jointly select the donors and define the introduction and elite crosses to ensure an overall consistency of genetic base broadening accounting for the performance and diversity available in both bridging and breeding populations.

### Genetic resources and pre-breeding

Different sources of diversity can be considered by commercial breeders. The most original, but which show a large performance gap with elites, are landraces (e.g. DH libraries derived from landraces, Strigens *et al.* 2013; Melchinger *et al.* 2017; Böhm *et al.* 2017) and first varieties derived from landraces. Since breeding industry is highly competitive, breeders are likely reluctant to introduce unselected genetic resources directly into the breeding germplasm despite they might carry favorable adaptation alleles to face climatic changes (McCouch *et al.* 2013; Hellin *et al.* 2014; Böhm *et al.* 2017). Instead, commercial breeders will prefer to consider elite inbred lines from other than their own program (Kannenberg 2001).

In this study, the external breeding program was designed to release every generation several improved lines, later considered as donors for genetic base broadening of the commercial breeding program. The external program started from a broader genetic diversity than the commercial program (on average, He = 0.283 compared to He = 0.133 at the end of burn-in) and was designed to maintain higher genetic diversity during selection (on average, He = 0.101 compared to He = 0.014 after sixty years). This was done to mimic in a simple way the outcome of the activity of several companies conducting separate programs and therefore maintaining a global diversity. The external program can also be viewed as a pre-breeding program since it aimed at improving genetic resources to reduce their performance gap with elites while maintaining genomewide diversity among the pre-breeding population (Figure 1). The situation where the commercial breeding program can access donors released twenty years ago mimicked the situation of private lines with expired plant protection act in maize (Mikel and Dudley 2006) or old public lines. The situation where the commercial breeding program can access donors released five years ago mimicked either donors released by pre-breeding programs (e.g. in maize the SeeD project, Gorjanc *et al.* 2016) or donors released by programs working a different genetic basis and targeting different environments (e.g. commercial varieties in inbred species accessible for breeding under the UPOV convention, Dutfield 2011).

The selection intensity was lower in the external breeding than in the commercial breeding programs (10% vs 5% of progeny selected, respectively). This was done to compensate the increased response to selection due to the higher genetic diversity and ensure that the donors released by the external program underperform the commercial breeding elites. It should be noted that donors outperforming elites might be encountered in practice when considering elite germplasm as source of diversity, but this situation was not considered in this study. In such a situation the direct introduction of donors would be clearly preferable.

Our results highlighted a clear beneficial effect of introducing external diversity in the elite program. This interest is all the more important as the level of introduced material is high, which highlights the importance of protection policies on long term genetic gain. More importantly, we show that the approach for introduction should be tuned given the type of external diversity that can be accessed (see next section).

### Interest of bridging relative to direct introductions in the elite pool

When considering recent and performing donors (five-year old), scenarios with introductions after bridging or direct introductions performed similarly. Conversely, for panel and twenty-year old donors, introductions after bridging yielded significantly higher mid- and long-term performance compared to direct introductions. Since donors (D) were less performing than elites, the fraction of progeny selected in donor by elite bridging families (DE progeny) carried on expectation less than half of donor’s genome (Allier *et al.* 2019b). Thus, progeny of introduction crosses after bridging (DExE) carried on expectation less than one fourth of the donor (D) genome. This D fraction includes favorable alleles but also unfavorable alleles brought by linkage drag, which number depends on the donor considered. Introductions penalized the mean breeding population performance in the first generations (Figure 3A-B). Next generations of recombination and selection partially broke the linkage between favorable and unfavorable alleles in introduced regions, resulting in a higher genetic gain than in the benchmark (Figure 3A-B) and an increase of the frequency of novel favorable alleles (Figure 3C). The more performing the donor, the less unfavorable alleles linked to favorable alleles and the more rapidly novel favorable alleles were introduced and spread in the breeding population (Figure 3C). In absence of bridging, the introduction progeny (DxE) carried on expectation one half of the donor genome. Consequently, the penalty due to introductions was more important and the conversion of additional diversity into genetic gain required more recombination events, i.e. recycling generations (Figure 3A-B). For panel donors showing a large performance gap with elites, the direct introductions were not converted into genetic performance. The high inter-family additive variance in this scenario (Figure S1 A) reflected the structuration of the breeding population into badly performing introduction families and performing elite families with only limited gene flow between them. Such behavior might be corrected by adding a constraint to force the recycling of introduction progeny in Eq. 1 when donors are too badly performing, which requires further investigations.

### Practical implementation in breeding programs

We considered a commercial breeding program with a genetic diversity at the end of the burn-in matching that of an experimental program reported by Allier *et al.* (2019a). Breeding programs ongoing for different species and breeders may present a diversity superior or inferior to the one that was simulated, which would make the importance of introductions lower or stronger than in the simulated scenarios, respectively. UCPC based OCS for genetic base broadening requires to genotype the candidate parents, including breeding material and potential donors, a genetic map and reliable marker effect estimates. This information is available in breeding programs that have already implemented genomic selection. In this study, we assumed fully homozygous inbred lines but considering heterozygote parents in UCPC based OCS is straightforward following the extension of UCPC to four-way crosses (Allier *et al.* 2019b). So similar approach could be tested for perennial plants or animal breeding schemes.

In scenarios with bridging, we considered by default two distinct bridging and breeding GS models. The prediction of elite (ExE) and introduction (DExE) crosses usefulness and the prediction within crosses were based on a model trained on the breeding progeny of the three corresponding previous generations. Considering a unique genomic selection model trained on both bridging and breeding progeny increased the prediction accuracy within introduction families (DExE) (Table 2). This higher selection accuracy favored the spreading of the introduced favorable alleles in the breeding population and resulted in an increased mid- and long-term performance (Figure 5). Furthermore, compared to use two distinct TS, a single TS led to introduce more bridging progeny (DE) for scenarios considering good performing donors (five-years old) and less for scenarios considering bad performing donors (twenty-years old) (Figure S3 A). Also, as we likely selected more accurately the introduction crosses (DExE) with a single TS, there was an increase in the proportion of those that contributed to the ten best lines, especially for twenty-year old and panel donors (Figure S3 B).

It is well known that the prediction accuracy is increased for larger TS (Hickey *et al.* 2014). At constant TS size, increasing the proportion of bridging progeny (DE) up to one third in the TS significantly increased the family variance prediction accuracy 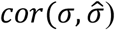 and within family prediction accuracy (*cor*(*u, û*)) in introduction families (DExE). Conversely, these higher proportions of bridging progeny (DE) in the TS significantly decreased 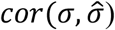 and *cor*(*u, û*) in elite families (ExE). The optimal balance between introduction and elite family prediction accuracies is likely data dependent as observed when considering genotypes and phenotypes simulated in different generations (Figure S4). For instance, considering later generations, a large proportion of DE in the TS penalized less the within elite prediction accuracy (Figure S4 C). The reason being that later breeding generations get closer to the external program germplasm (Figure 4). The optimal balance between bridging and breeding progeny in the training set might be defined using an optimization criterion such as the CDmean (Rincent *et al.* 2012) extended to account for linkage disequilibrium as suggested by Mangin *et al.* (2019).

We proposed to implement bridging at constant cost by splitting the breeding population into a small bridging population and a large breeding population. This involves practical changes in the breeding organization that remain to be studied. We considered equal family sizes and within family selection intensities for bridging and breeding families. However, in practice different within family selection intensities can be considered in UCPC based OCS (File S2) and one may want to modulate the selection intensity among families, e.g. select less intensively in bridging and more intensively in breeding families. We could consider the selection intensities as fixed parameters regarding breeding objectives or as variable parameters to be optimized. The effect and the optimization of within family intensities in bridging and breeding requires further investigations. We considered a selection accuracy *h* = 1 for cross selection, for sake of facility. However, we observed that within family prediction accuracies were variable (Table 2, Figure 6). Note that *a priori* within family accuracy can be accounted for in UCPC based OCS (File S2). For instance it would give less importance to predicted variance for crosses with *a priori* low within family accuracy. The consequences on short- and long-term UCPC based OCS efficiency need to be investigated. In bridging, we gave more importance to performance than to diversity (*α* = 0.7) when selecting bridging crosses in order to reduce the performance gap between donors derived material and elites. When giving less weight to the performance than to the diversity, i.e. *α* = 0.3, we observed non-significant changes on the short- or long-term performance for scenarios with five-year and twenty-year old donors and a significant increase of long-term performance and novel favorable allele frequency for the scenario with panel donors (Figure S2 A-C). This suggested that for unimproved donors, selecting too strongly for performance in bridging favors the first elite recipient genome contribution and limits the introduction of novel favorable alleles. Further investigations are required to better define this parameter for practical implementation.

### Outlooks

We considered an inbred line breeding program corresponding to selecting lines on *per se* values for line variety development or on testcross values with fixed tester lines from the opposite heterotic pool for hybrid breeding. In this case, the use of testcross effects estimated on hybrids between candidate lines and tester lines is straightforward. The extension to hybrid reciprocal breeding is of interest for genetic broadening in several species such as maize and hybrid wheat (Longin and Reif 2014). In this context it is possible to account for the complementarity between heterotic groups in UCPC based OCS to complementarily enrich and improve both pools. This would require to include dominance effects in UCPC based OCS. We considered a single trait selected in both the external and the commercial breeding programs in the same population of environments for a total of eighty years. These assumptions should be relaxed in further simulations. Firstly, it is well recognized that genetic resources suffer agronomic flaws (e.g. lodging, Tallury and Goodman 2001; Longin and Reif 2014) or miss adaptation (e.g. flowering time) that should be accounted for during pre-breeding and introduction in breeding. In such a multi-trait context, the multi-objective optimization framework proposed in Akdemir *et al.* (2019) can be adapted to UCPC based OCS. Secondly, in practice several public pre-breeding programs or competitor programs can be considered as sources of candidate donors for genetic base broadening. These programs likely did not select for the same target environments and are themselves continuously enriched in new allelic variation. Thirdly, in a context of climate change and rapid evolving agricultural practices, breeding targets are expected to change (e.g. emerging biotic or abiotic stresses). Considering a more realistic context, where donors are released by different programs selecting in different environments and for different traits changing over time, likely makes the interest of maintaining genomewide genetic diversity through genetic base broadening even more important than highlighted in this study.

## AUTHOR CONTRIBUTION STATEMENT

AC, ST, CL and LM supervised the study. AA and ST worked on the simulator. AA performed the simulations, analysis and wrote the early version of the manuscript. All authors reviewed and approved the manuscript.

## ACKNOWLEDGMENTS

This research was funded by RAGT2n and the ANRT CIFRE Grant n° 2016/1281 for AA. It beneficiated from a financial support and helpful discussions with the members of the “Gdiv-Selgen” and “R2D2” projects within the framework of the INRA “Selgen” meta-program.

